## Supplementary material for "Characterization of genetic and molecular tools for studying the endogenous expression of *Lactate dehydrogenase* in *Drosophila melanogaster*": Figure S1

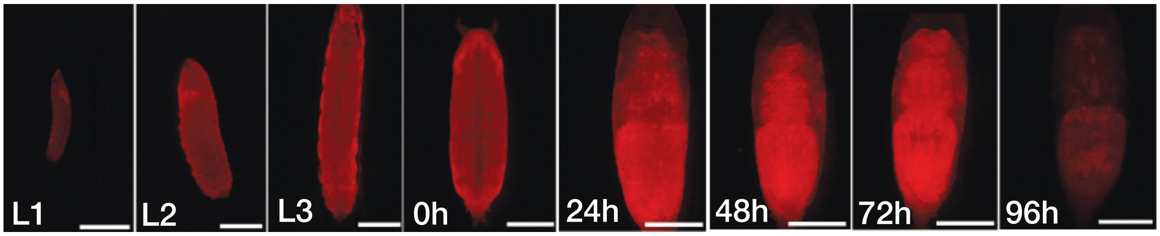


**Figure S1. Expression of *Ldh-mCherry*$^{Genomic}$ during larval development.** The *Ldh-mCherry*$^{Genomic}$ spatial expression pattern is consistent with previous studies, with Ldh-mCherry$^{Genomic}$ being expressed at high levels in (A) the body wall muscle. However, unlike Ldh-GFP$^{Genomic}$, the (A) expression of mCherry$^{Genomic}$ fusion protein persists throughout much of pupal development (compare with Figure 1B).
