## Supplementary material for "Characterization of genetic and molecular tools for studying the endogenous expression of *Lactate dehydrogenase* in *Drosophila melanogaster*": Figure S2

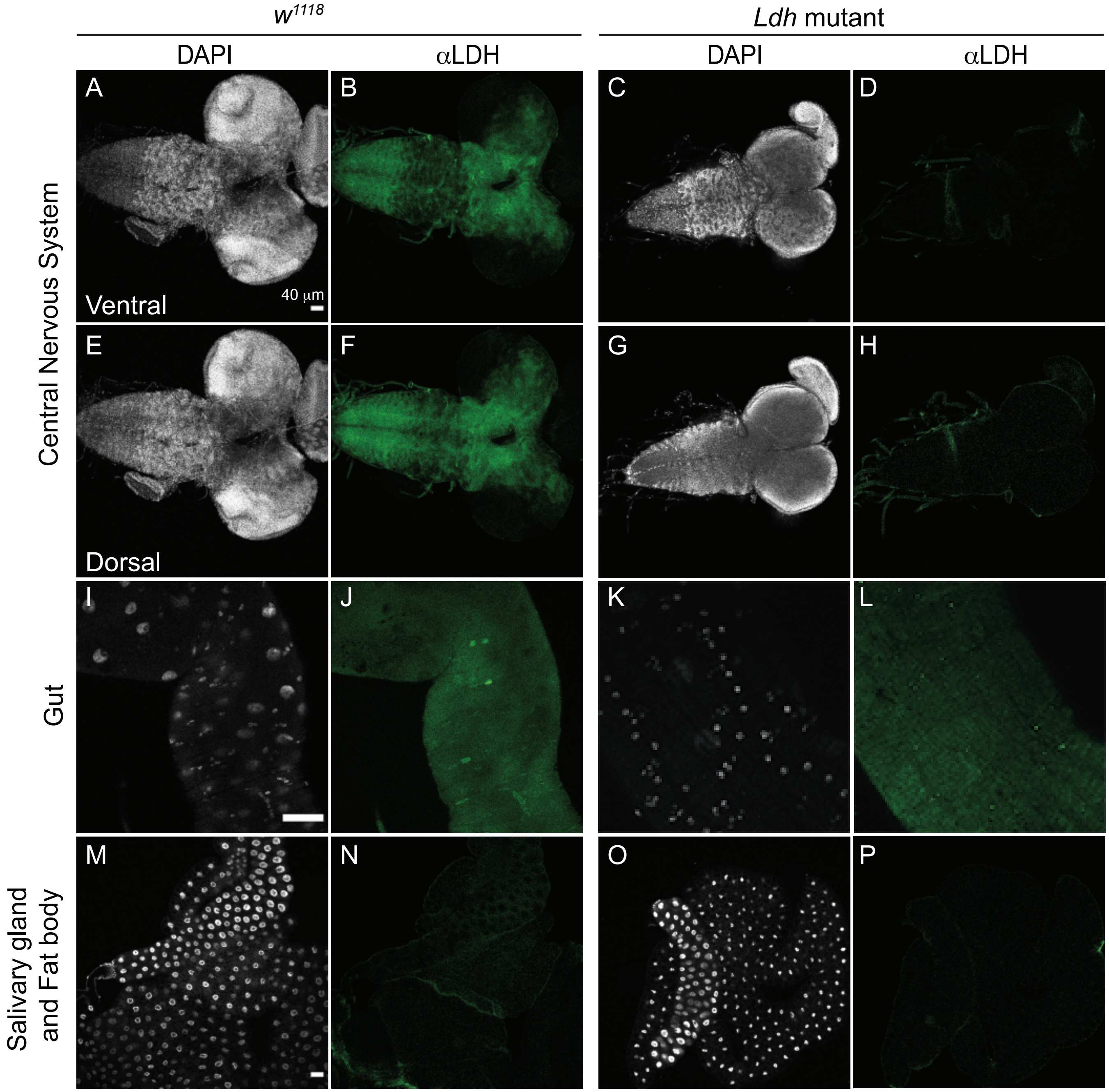


**Figure S2. αLdh immunostaining in control and *Ldh* mutant larval tissues.** Representative confocal images of the (A-H) central nervous system, (E-H) intestine, (I-L) fat body, and (M-P) salivary gland dissected from third instar $w^{1118}$ and $Ldh^{16}$ larvae. DAPI is shown in white and *Ldh* expression is shown in green. Note the brightness in (L) was increased to highlight the lack of staining in small cells within AMP clusters. The scale bar in all images represents 40 μM. The scale bar in (A) applies to panels (B-H), the scale bar in (I) applies to panels (J-L), and the scale bar in (M) applies to panels (N-P).
